# Model-based inference of cell cycle dynamics captures alterations of the DNA replication programme

**DOI:** 10.1101/2025.03.19.644216

**Authors:** Adolfo Alsina, Marco Fumasoni, Pablo Sartori

**Affiliations:** Instituto Gulbenkian de Ciência, Rua da Quinta Grande 6, 2780-156 Oeiras, Portugal

## Abstract

The cell cycle of eukaryotic cells consists of several processes that must be carefully orchestrated and completed in a timely manner. Alterations in cell cycle dynamics have been linked to the onset of various diseases, underscoring the need for quantitative methods to analyze cell cycle progression. Here, using a combination of high-throughput experimental data and theoretical modeling, we develop RepliFlow, a model-based approach to infer cell cycle dynamics from flow cytometry data of DNA content in asynchronous cell populations. We model the DNA content distribution as the result of both noisy single-cell measurements and the population’s age structure. We show that RepliFlow captures not only changes in the length of each cell cycle phase but also alterations in the underlying DNA replication dynamics. RepliFlow is species-agnostic and recapitulates results from more sophisticated analyses based on nucleotide incorporation. Finally, we propose a minimal DNA replication model that enables to derive microscopic insights about origin firing rates and replication fork speed from population-wide DNA content measurements. Our work presents a scalable framework for inferring cell cycle dynamics from flow cytometry data, enabling the characterization of replication program alterations.

---

The cell cycle is the series of events between successive divisions, requiring precise coordination of molecular duplication to ensure viability. Disruptions in this process can impair division and lead to genetic instability and cell death [1–3].

Most eukaryotic cell cycles have a highly conserved structure, being composed of four major phases [4]. During the first phase G_1_ (Gap 1), cells prepare for DNA replication. In the subsequent S phase (synthesis) the DNA is replicated. Finally, cells enter G_2_ (Gap 2) phase, which paves the way for M (mitosis), when the cell physically divides. A deep understanding of the constraints regulating the cell cycle requires the development of methods that can accurately quantify the timing of its phases across a wide spectrum of conditions.

Microscopy-based fluorophore reporters have been developed to track cell cycle dynamics at the single-cell level in different organisms [5–7]. While powerful, these methods are limited in scalability due to genetic requirements and lengthy acquisition times. A compelling alternative is represented by the use of DNA content as a proxy for cell cycle progression. With a single dye intercalating the DNA, and without the need for genetic manipulations, the DNA content of thousands of single cells can be readily measured using flow cytometry [8, 9]. As the DNA content changes in a stereotypical way as cells progress through the cell cycle, this method allows in principle to distinguish relative abundance of three out of the four main cell cycle phases (G_1_, S and G_2_*/*M) in parallel, in hundreds of different conditions.

However, current approaches to infer cell cycle dynamics from the information present in the distribution of DNA content across an asynchronous population (DNA profiles) are not grounded on biophysical models of cell cycle regulation [10–12] and they commonly rely on heuristic definitions of the cell cycle phases. Here, we develop RepliFlow, a new framework to infer cell cycle dynamics from flow cytometry data based on a biophysical model of the cell cycle. Our approach is cell-type agnostic and depends only on a small number of interpretable parameters, encoding not only the time allocated to each cell cycle phase but also possible alterations of the DNA replication dynamics.

Using a combination of published datasets and targeted experimental perturbations of the cell cycle dynamics, we show that we can accurately infer parameters characterising the cell cycle dynamics in a large number of different conditions and cell types, requiring solely DNA content measurements. Furthermore, by introducing a microscopic model of the DNA replication dynamics, we also show that this approach can infer microscopic observables, such as the typical replication fork velocity, with an accuracy on par with modern sequencing technologies at a fraction of the cost and effort. Taken together, our framework provides a quantitative and scalable approach to study time allocation in growing cell populations, paving the way to future studies aimed at understanding the biophysical constraints underlying this process.

## RESULTS

### Inference of cell cycle dynamics recapitulates cell cycle time allocation in *S. Cerevisiae*

Consider a population of cells growing asynchronously. As each cell is in a different phase of its cycle (Figure 1(a)), the population is characterised by an empirical distribution of DNA abundance per cell, which depends on the relative duration of each phase. Here, we develop a general likelihood-based approach to infer the fraction of time allocated to each cell cycle phase from such distribution (see also Materials and Methods).

**FIG. 1.**
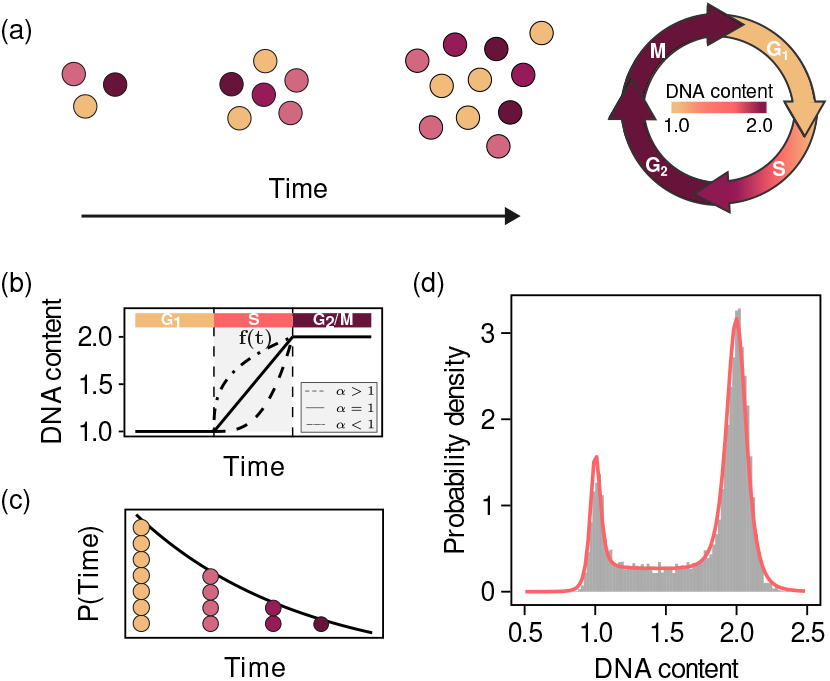
Distribution of DNA content as a combination of single-cell and population level processes. (a) Schematic of the time evolution of an exponentially growing asynchronous cell population. (b) Dynamics of DNA content during the length of the cell cycle. (c) Distribution of time along the cell cycle (age) in an exponentially growing asynchronous cell population. (d) Distribution of DNA content in an asynchronous cell population and resulting maximum likelihood fit of our model (red).

The centerpiece of our analysis is ***y*** = {*y*_1_, …, *y*_*N*_}, the empirical distribution of DNA content in a population of *N* cells. The distribution ***y*** depends on the cell cycle dynamics of individual cells, as well as on the age distribution of the population. The likelihood of measuring ***y*** = {*y*_1_, …, *y*_*N*_} can be written as

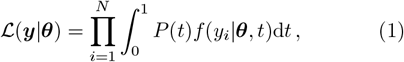

where *P* (*t*) represents the cell cycle age distribution across the population, *t* the time elapsed since the last division event; and *f* (*y* | ***θ***, *t*) is a function that quantifies the amount of DNA content in a cell of age *t* that is characterised by parameters ***θ***. The vector ***θ*** includes parameters such as the duration of the different cell cycle phases and also parameters that account for the effect of technical noise. For a given DNA content distribution ***y***, we infer the parameter values that maximise the log-likelihood function

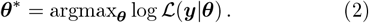

To completely determine the model, it remains to specify the forms of *P* (*t*) and *f* (*y*|***θ***, *t*). These contributions are schematically represented in Figure 1(b-c). First, we consider the measured amounts of DNA to result from the combination of deterministic single-cell dynamics and technical noise (see Material and Methods). We represent the deterministic dynamics of the amount of DNA content (Figure 1(b)) by a continuous piecewise function

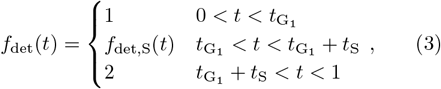

where *t*_*i*_ is the fraction of time allocated to phase *i* and *f*_det,S_ is a monotonically increasing function describing the change of DNA content during S phase. To account for technical noise, we write *f* (*y*| ***θ***, *t*) as a probability distribution with mean *µ* = *f*_det_(*t*) and variance *σ*^2^.

We model DNA replication dynamics during S phase using a phenomenological model that accounts for defects in early and late replication:

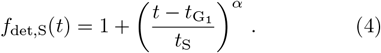

For *α* = 1 the model reduces to a linear model of DNA replication, whereas *α <* 1 and *α >* 1 correspond, respectively, to defects in early and late replication (Figure 1(b)).

Next, we focus on the cell cycle age distribution *P* (*t*) (Figure 1(c)). The distribution of ages in a dividing population undergoing exponential growth is also exponential [13]. Specifically, for a homogeneous population of cells, the age distribution can be written as

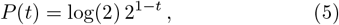

where *t* is the elapsed fraction of time since the start of the cell cycle.

Equation 1 together with the definitions of *f* (*y* |***θ***, *t*) and *P* (*t*) constitute a closed optimization problem on the parameters ***θ***. Given an empirical distribution of DNA content, this approach infers the allocated time fractions and technical noise parameters that maximise the likelihood function.

To validate our model, we tested it on the DNA profile of an exponentially growing asynchronous *S. cerevisiae* population (Figure 1(d)), correctly reproducing the main features of the data. For the DNA distribution of Figure 1(d), obtained from a population grown in media supplemented with 2% glucose at 30^*°*^*C*, our method estimates that cells allocate 9.7% of time to G_1_ phase, 23.3% of time to S phase and 67% of time to G_2_*/*M. For the measured doubling time in these conditions of 117.2 min, these fractions correspond to absolute durations of 11.37 min, 27.30 min and 78.52 respectively. The inferred phase durations qualitatively align with the expected time spent in each phase under these conditions. Regarding replication dynamics, for this profile we find a value of *α* = 0.96, indicating that the replication dynamics are well described by a constant replication speed throughout S phase, reflected in a linear increase of DNA content.

Thus, our approach provides a new way of estimating cell cycle dynamics that is fast, robust and accurate.

### Analysis of the haploid yeast deletion collection captures alterations in the cell cycle

Having established our inference framework, we set out to show that it can accurately capture variability in the time allocated to the different cell cycle phases. With that goal in mind, we resort to analysing the FACS data of the haploid yeast deletion collection, a comprehensive dataset of all non-essential single-gene deletion strains in haploid *S. cerevisiae* [14, 15]. As several mutants in the deletion collection impact key cell cycle processes, it provides a suitable testing ground to reveal how cell cycle time is allocated in budding yeast mutants.

The analysis of the deletion collection reveals large variablity in the relative time allocated to all three distinguishable phases across mutants (Figure 2(a)). Furthermore, the distribution of fitting scores (Figure 2(b), see Material and Methods for details) shows that 95% of the deletion collection is fitted with a high degree of accuracy *(s >* 0.95). We find a strong negative correlation between the percentage of time spent in G_1_ and G_2_/M phases, as previously reported [16]. This correlation accounts for most of the variation between individual mutants, indicating that frequently an extension of one of the gap phases comes at the expense of the length of the other (Supplementary Figure). As the experimental design used to obtain this dataset is prone to presenting technical differences between plates, in the following we normalise our results by the in-plate controls (z-scores, see Materials and Methods for details).

**FIG. 2.**
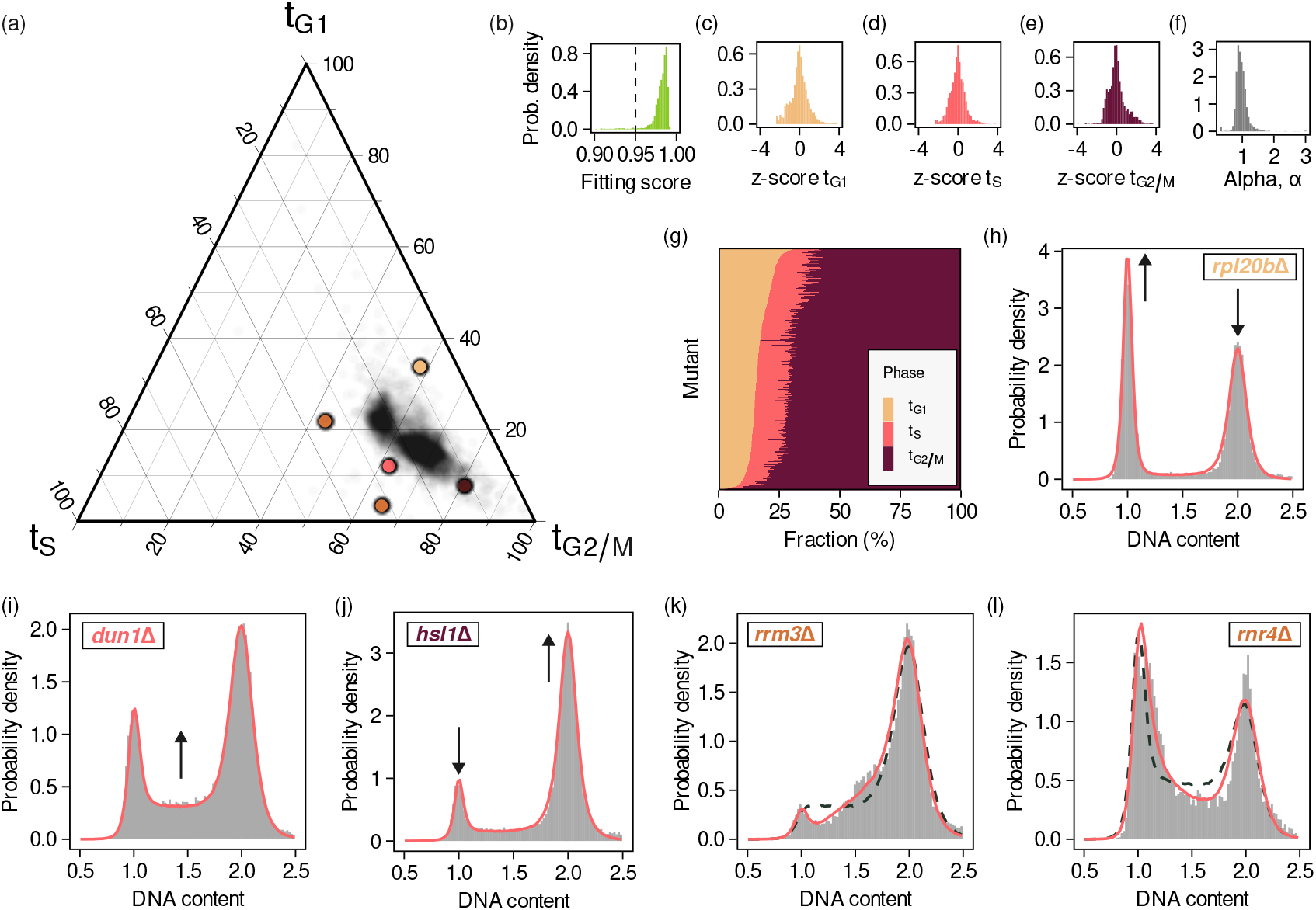
Analysis of the yeast deletion collection recovers extended cell cycle phase durations and alterations of the replication programme. (a) Ternary plot of the fraction of time spent in each cell cycle phase. (b) Distribution of fitting scores of the deletion collection profiles. The dashed line corresponds to the minimum fitting score required for all subsequent analysis. (c-e) Distribution of z-scores of the fraction of time spent in G1 (c), S (d) and G2/M (e) phase. (f) Distribution of alpha values of the deletion collection profiles. (g) Heatmap of the fraction of time allocated to G1, S and G2/M for all deletion collection mutants. (h-j) Example profiles and fits of mutants with extended G1 (f), S (g) and G2/M phase (h). Example profiles and fits of mutants with fitted *α* values smaller (k) and larger (l) than one.

The inferred parameter distributions (Figure 2(c-f)) are peaked around 0, indicating that the bulk of the deletion collection can be well described by the average time fractions allocated to the different phases across the whole collection together with a linear model of DNA replication (Figure 2(f)). Conversely, the tails of the z-score distributions correspond to mutants with shortened or extended phase durations whereas the tails of the *α* distribution correspond to mutants with defects during early or late DNA replication, respectively. The homogeneity of the bulk of the deletion collection is also reflected in Figure 2(g), where it can be seen how a large number of mutants display only subtle differences in the fraction of time allocated to each cell cycle phase.

To show that our method reliably captures known cell cycle alterations, in the following we focus on representative mutants from the tails of the parameter distributions (highlighted in Figure 2(a)) for which the biological mechanism underlying the corresponding cell cycle alteration is well known.

First, we briefly focus on mutants presenting extended duration of one cell cycle phase. We identify *rpl20b*Δ mutants as having an extended G_1_ phase (Figure 2(h)). Deletion of this ribosomal subunit-encoding gene reduces protein production, prolonging G_1_ as cells grow to the appropiate size for S phase entry [17]. The extension of S phase is reflected in the profiles of *dun1*Δ mutants (Figure 2(i)), lacking a protein kinase involved in production of deoxyribonucleotides, and prolonging the time needed to complete genome replication [18– Finally, mutants lacking *HSL1* (Figure 2(j)), a gene involved in cytokinesis, exhibited extended G_2_*/*M duration [21]. Taken together, our results show that RepliFlow systematically captures variability in the time allocated to each cell cycle phase across mutants, recapitulating previously known cell cycle alterations.

Moreover, our approach not only captures lengthenings of the duration of each cell cycle phase but also alterations of the DNA replication dynamics. As defects during early or late DNA replication would be reflected in the inferred values of the parameter *α*, we next focus on the tails of the *α* distribution.

Figure 2(k) shows the DNA profile of a *rrm3*Δ mutant strain. *RRM3* encodes for a DNA helicase, crucial for the removal of protein obstacles to the DNA replication forks [22–24]. In *rrm3*Δ mutants stalled forks delay the completion of DNA replication, leading to the observed build-up of cells with incompletely duplicated genomes at the end of S phase. We infer a value of the parameter *α* = 0.48 *<* 1 (solid line, dashed corresponds to a linear model), reflecting the observed accumulation of cells close to the G_2_/M peak and corresponding to the presence of defects during late DNA replication.

The profile of a *rnr4*Δ strain is shown in Figure 2(l). *RNR4* catalyzes the synthesis of dNTPs [25, 26], the required building blocks for DNA replication, causing problems initiating DNA replication in mutants lacking this gene. The presence of defects during early replication is reflected in the inferred value of *α* = 1.68.

In summary, we have introduced a model that correctly captures lengthenings of the different cell cycle phases and alterations in the DNA replication programme. These alterations are characterised using a minimal, uniparametric model of DNA content changes along the length of the cell cycle. Together, the proposed approach permits the identification of strains associated with specific cell cycle alterations from the tails of the parameter distributions.

### Quantifying graded perturbations of the cell cycle dynamics

Gene deletions are powerful tools to perturb cellular processes. However, deletions are restricted to non-essential genes and limited by the all-or-nothing nature of the perturbations they produce. To study the response to a graded perturbation, we performed experiments where cells are exposed to methyl methanesulfonate (MMS), an alkylating agent that modifies bases and perturbs DNA replication [27, 28]. To quantify the impact of MMS of the DNA replication dynamics, we systematically vary the concentration of the chemical across more than two orders of magnitude.

We find that for small concentrations of MMS, the DNA profile of MMS-treated cells resembles that of untreated cells (Supplementary Material). However, increasing the MMS concentration causes the profiles to become increasingly skewed (Figure 3(a)), with an increase in the number of cells close to the G_2_*/*M peak, as a result of cells stalling before completing DNA replication.

**FIG. 3.**
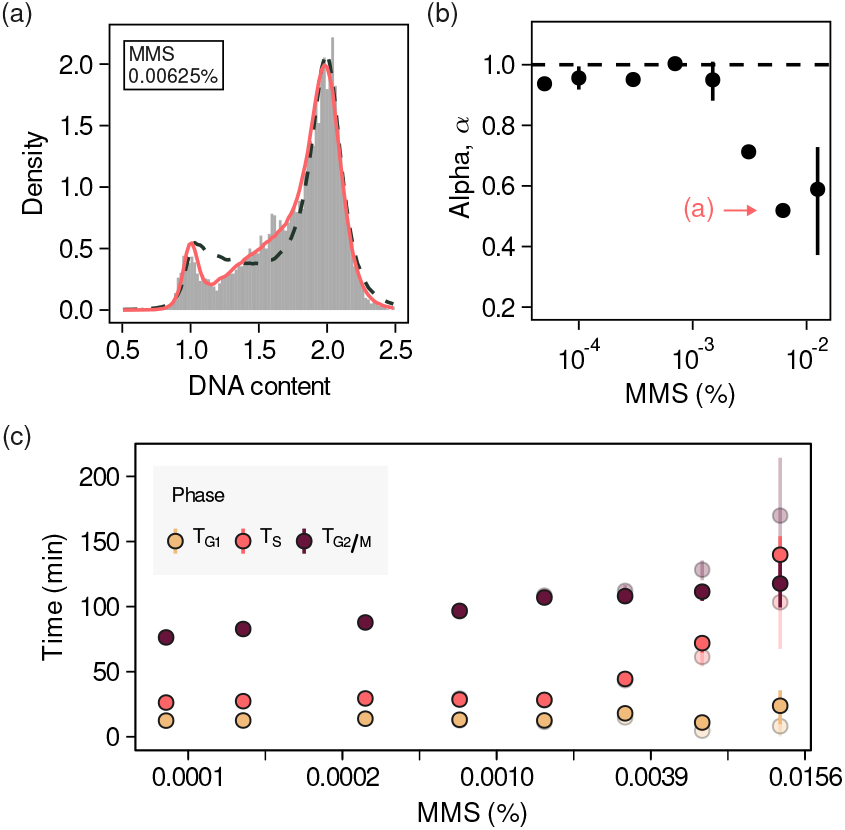
Quantifying a graded perturbation of the DNA replication dynamics. (a) DNA profile of cells grown in the presence of MMS (0.00625%) presenting defects in late DNA replication. Solid lines correspond to fits of the *α* model to the data. Dashed lines correspond to fits of a linear model.(b) Fitting parameter *α* as a function of the MMS concentration in the media. Points correspond to the average across four independent experiments and bars correspond to 95% confidence intervals. The MMS concentration corresponding to the profile shown in (a) is highlighted. (c) Time allocated to each cell cycle phase as a function of the MMS concentration inferred using a linear (transparent) and alpha (solid) replication models.

In Figure 3(b) we show the dependence of the parameter *α* on the concentration of MMS. As the MMS concentration is increased, we observe a decrease in the inferred values of *α*, corresponding to a shift from a linear DNA replication programme (*α* ≈ 1) at low doses to altered replication dynamics at high doses. The DNA replication dynamics at high MMS concentrations are dominated by the presence of late defects difficulting completion, as reflected in the inferred values of *α <* 1.

By independently measuring the doubling time of cells in each condition (Materials and Methods), we quantify how the time allocated to each cell cycle phase changes as a function of the MMS concentration (Figure 3(c)). The total amount of time spent in each cell cycle phase is defined as *T*_*i*_ = *T*_d_ *t*_*i*_, where *i* = {G_1_, S, G_2_*/*M}, *T*_d_ is the doubling time of cells in these conditions and *t*_*i*_ is the fraction of time spent in phase *i*. We find that for large MMS concentrations the duration of all cell cycle phases is extended, with the most prominent increase corresponding to the time spent in S phase.

Our results are compatible with an interpretation where for low MMS concentrations the induced damage can be efficiently overcome, not impacting the overall cell cycle dynamics. However, the damage induced by larger concentrations of MMS requires cell to spend more time in S phase in order to be repaired and DNA replication to be completed. The transition between these two regimes can be readily identified thanks to the quantitative character of this approach.

### Characterisation of mammalian replication dynamics from DNA content alone

Although in previous sections we have focused on budding yeast datasets, the model is based on the conserved structure of the eukaryotic cell cycle. To illustrate the broader applicability of this framework, we next show that we can also characterise cell cycle dynamics in mammalian cell lines.

A common approach to studying DNA replication in mammalian cells involves nucleotide incorporation assays [29–31]. Fluorescent analogs such as 5-Ethynyl-2’-deoxyuridine (EdU) integrate into newly synthesized DNA (Figure 4(a)), with EdU signal intensity reflecting replication rate and the number of actively replicating cells. Commonly, the fraction of cells in each cell cycle phase is calculated using a gating approach based on an ad-hoc geometric definition of the various cell cycle phases in the 2D DNA content-EdU plane (Figure 4(b)).

**FIG. 4.**
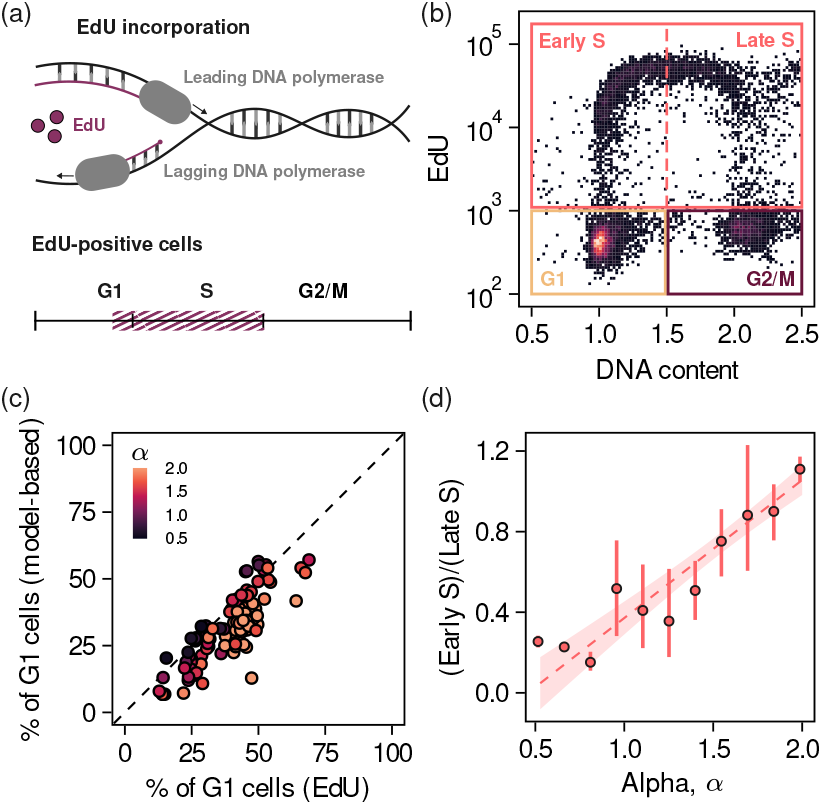
Replication dynamics of mammalian cells obtained from DNA content alone. (a) Schematic of an EdU incorporation experiment. (b) 2D histogram of EdU intensity and DNA content (data from Rainey et al., *Cell Reports*, 2020). (c) Fraction of cells in G1 phase estimated from the EdU-DNA content histograms vs estimated from the DNA profiles using our method (d) Ratio of the fraction of cells in early and late S phase estimated from the EdU-DNA content histograms as a function of the fitting parameter *α*.

We applied our framework to the EdU incorporation experiments of [32], where the authors screened mutants with altered response to inhibitors of DNA replication initiation. First, we calculated the fraction of cells in each cell cycle phase using RepliFlow (Materials and Methods). Next, we calculated these same quantities starting from ad-hoc regions defined in the EdU-DNA content plane and compared the fraction of cells in G_1_ phase obtained using both approaches. In Figure 4(c)) we show that we reliably estimate the fraction of cells in G_1_ phase. We chose the fraction of cells in G_1_ as this is the only phase that can be unambiguously identified using the gating approach across all conditions (Supplementary Material). The strong correlation between the estimates obtained by the proposed method and the gating approach shows that we can indeed estimate the fraction of cells in different cell cycle phases from DNA content alone in a wide range of conditions, including pathological ones that strongly disturb cell cycle progression.

More importantly, we correctly characterise the DNA replication dynamics, as illustrated by the correlation between the ratio of the fraction of cells in early and late S phase calculated using the gating approach and the fitting parameter *α* (Figure *3(d)). Our results further show that α* serves as a proxy for the type of DNA replication defects, as it accurately captures alterations in the replication programme, in agreement with the results of the gating approach. Thus, the proposed method offers a robust alternative to quantify replication dynamics from DNA content alone consistent with more costly and time consuming experimental approaches.

### Microscopic model of the replication dynamics

RepliFlow relies on the specification of *f*_det_(*t*), the change in DNA content over the course of the cell cycle. The change in DNA content is specified by the dynamics of DNA replication, which has been extensively studied from a theoretical point of view in eukaryotes [34–39]. To illustrate the modular nature of our framework, we propose a minimal model of DNA replication that does not take into account the spatial configuration of DNA and show that it can accurately infer microscopic properties of the replication dynamics from DNA content measurements.

Figure 5(a) schematically illustrates the main components of the microscopic model. In brief, DNA replication starts from specific locations along the genome called origins of replication. During replication phase, licensed origins fire with rate *γ*. Upon firing, each origin recruits two replication forks that travel the genome in opposite directions (left) at speed *v*. When two replication forks meet, the DNA between their corresponding origins has been fully replicated and they detach from the DNA (center). As origin firing is a stochastic process, some origins are passively replicated and rendered inactive for the reminder of the cycle (right).

**FIG. 5.**
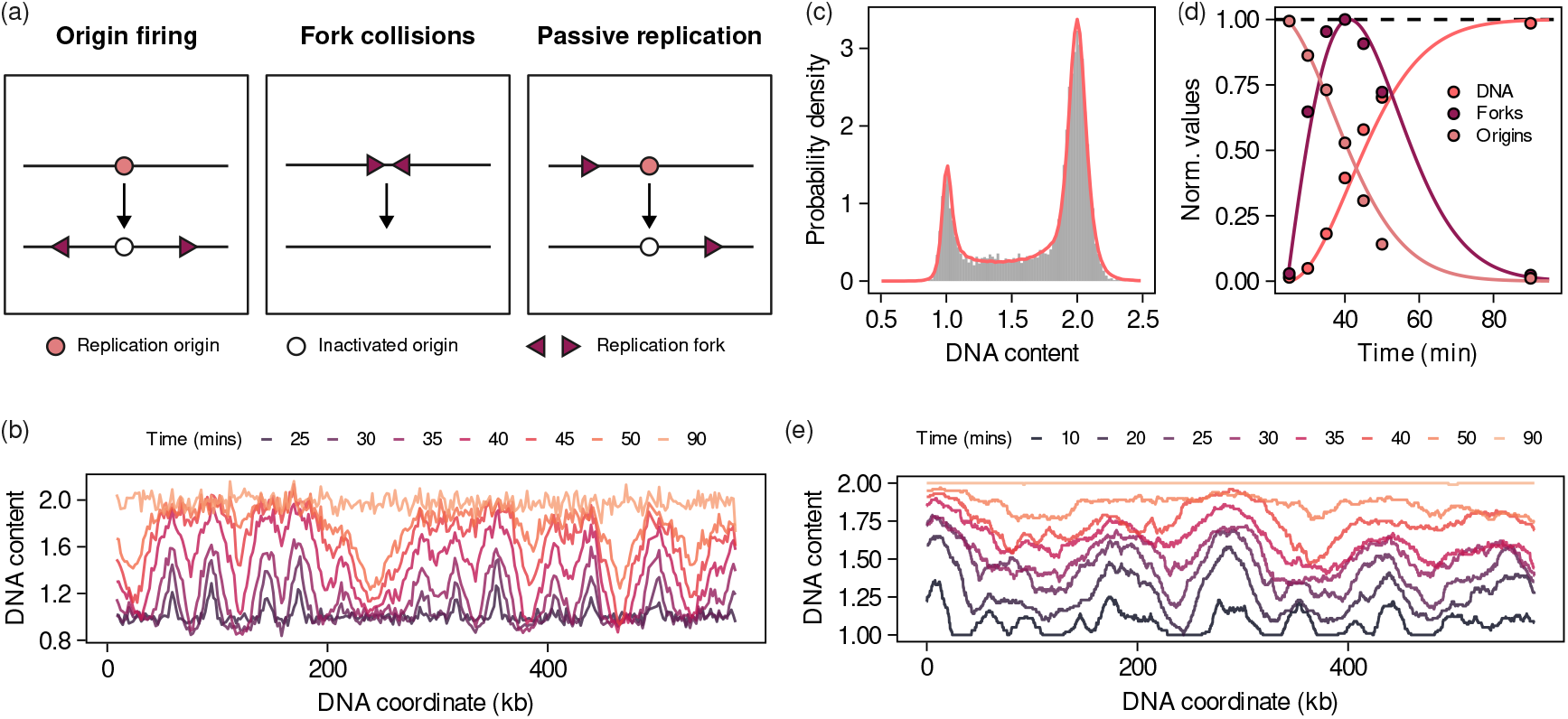
A minimal model of DNA replication quantitatively reproduces microscopic replication dynamics. (a) Cartoon of the processes included in our phenomenological microscopic DNA replication model. (b) DNA sequencing data of chromosome V of synchronised populations of budding yeast. Data taken from Müller et al., *Nucleic acids research*, 2014. [33] DNA profile and fit of the data shown in Fig. 1(d) using our microscopic DNA replication model. (d) Comparison of the microscopic model predictions against data from Müller et al., *Nucleic acids research*, 2014. [33] The solid lines correspond to a simulation of the coarse-grained model calibrated using data from Fig. 1(d). Points correspond to data extracted from the deep sequencing time course (Fig. 5(b)). (e) Simulated dynamics of DNA content in chromosome V of budding yeast using the reported origin locations in this chromosome and the microscopic parameters inferred from (c).

**FIG. 6.**
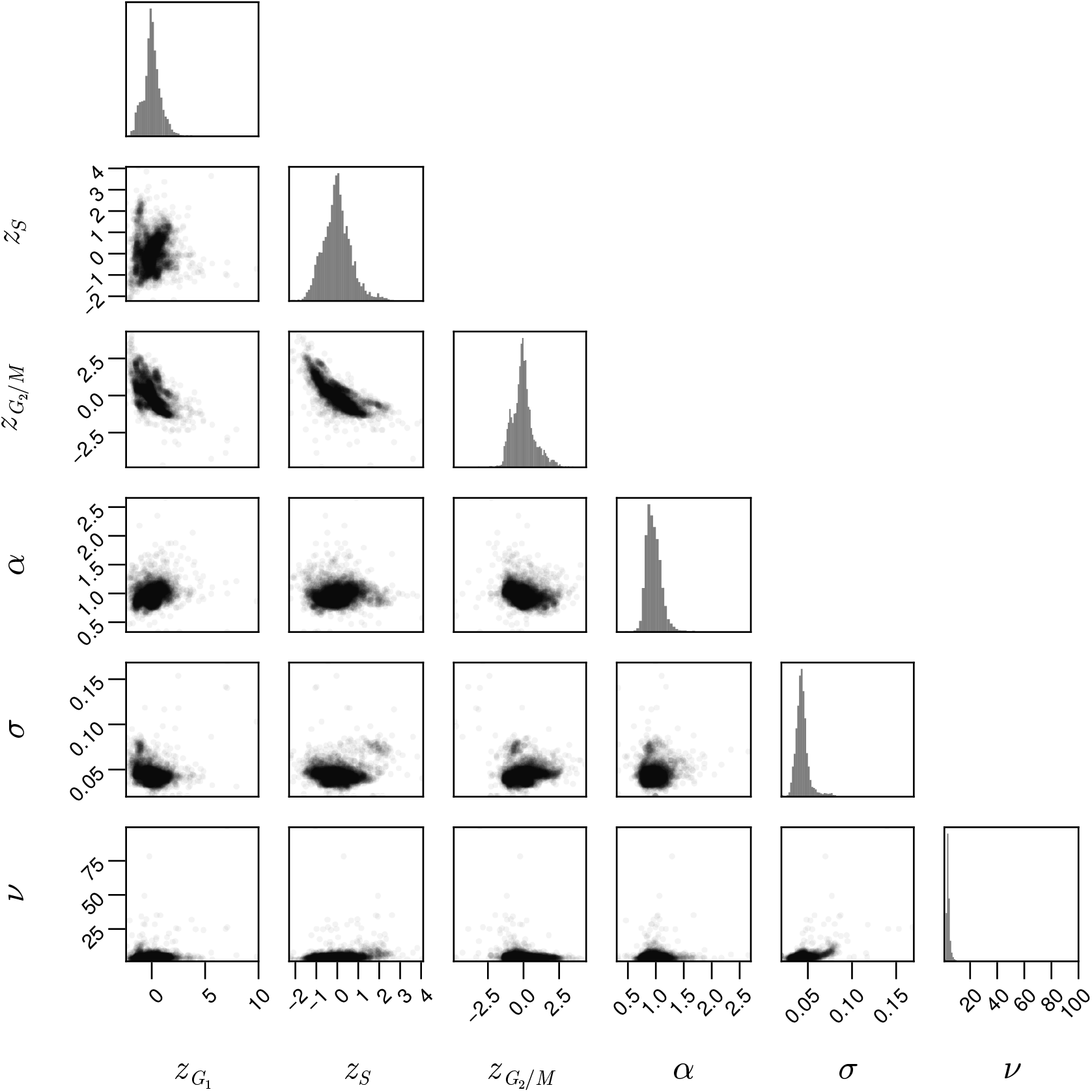
Supplementary figure. Paired plot depicting correlations between the inferred parameters for the yeast deletion collection.

**FIG. 7.**
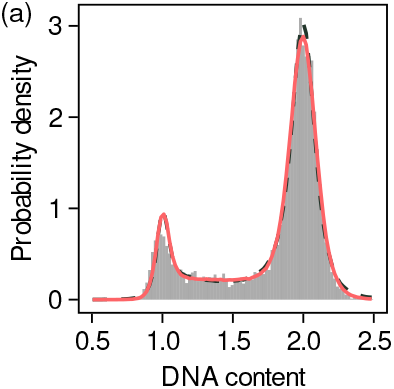
Supplementary figure. (a) DNA profile of cells grown in the presence of MMS (0.0015%). Solid lines correspond to fits of the *α* model to the data. Dashed lines correspond to fits of a linear model.

**FIG. 8.**
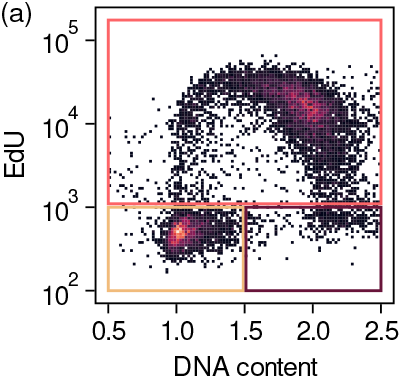
Supplementary figure. (a) 2D histogram of EdU intensity and DNA content (data from Rainey et al., *Cell Reports*, 2020) for which the identification of cells in S and G2/M cells is ambiguous.

We propose a mean-field microscopic model of DNA replication describing the temporal evolution of the number of active forks over time *n*, the number of licensed origins *ω* and the fraction of replicated DNA *ϕ*

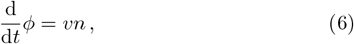

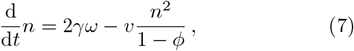

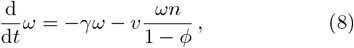

where *v* is the fork velocity and *γ* the origin firing rate. A detailed derivation of this model can be found in the Materials and Methods section.

To test whether RepliFlow together with the microscopic model can accurately infer microscopic parameters from DNA content profiles we turn to the synchronous deep sequencing experiments from Müller et al. [33] (Figure 5(b)). The high temporal and spatial resolution of these experiments makes them a suitable baseline against which to test the performance of the model. We extract from the DNA sequencing data the time series of the relevant quantities for the microscopic model: the fraction of replicated DNA, the number of licensed origins and the number of active forks (see details in the Materials and Methods).

To test the model, we first infer the microscopic parameters from DNA content measurements. We replace *f*_det,S_ in Equation 3 by *ϕ* and expand the parameter set ***θ*** so it now contains the microscopic parameters (*v, γ*). In Figure 5(c) we show the fit to a WT DNA profile. We obtain parameter estimates *v* = 1.76 kb*/*min and *γ* = 0.015 min^*−*1^, consistent with previous reports of replication fork velocity in budding yeast [40].

Using these estimates for the microscopic parameters, in Figure 5(d) we show that the microscopic model accurately reproduces the dynamics of the different observables extracted from the sequencing experiments of [33]. Importantly, in this comparison the microscopic model has no free parameters, as the fork velocity and origin firing rate are inferred from the DNA profile in Figure 5(c). Therefore, fitting the microscopic model to FACS data we can predict the dynamics of deep sequencing experiments. In Figure 5(e) we show a simulated time series of the DNA content distribution for chromosome V of budding yeast during synchronous replication (see Materials and Methods for details). Our results reveal that some spatial features can be reproduced starting from the reported origin locations and the microscopic parameters inferred using the model.

In summary, we have shown that a minimal DNA replication model depending only on two parameters can provide quantitative insights into the microscopic workings of DNA replication at a fraction of the cost, effort and time of sophisticated sequencing technologies.

## DISCUSSION

The study of cell cycle dynamics has long benefited from a wealth of available experimental techniques that allow to measure cell cycle progression both at the molecular [41, 42] and the population levels [27]. To extract maximum information from large datasets, new theoretical and computational tools that leverage our knowledge about cell cycle dynamics are required. Here, we have introduced RepliFlow, a new method to infer cell cycle dynamics from flow cytometry data based on a biophysical model of DNA content changes during cell cycle progression.

This approach extracts the duration of most cell cycle phases from DNA profiles of asynchronous cell populations, which can be obtained at a small cost and in a short time using a conventional flow cytometer, making our method particularly suitable for high-throughput analysis. We characterise DNA profiles in terms of the combination of single-cell and population level contribu-tions. At the single-cell level, we introduce a biophysical model of DNA content dynamics during the length of the cell cycle. Together with the age structure of the population and the effect of technical noise, these contributions describe the experimentally measured DNA profiles.

The method not only has less parameters than common alternatives, but they are interpretable, corresponding to the duration of the main cell cycle phases and the dynamics of DNA replication. Furthermore, the model captures alterations in the replication dynamics encoded in a single parameter *α*, that permits us to distinguish defects in early and late DNA replication.

Using chemical perturbations of the cell cycle, we showed that a quantitative approach can identify the transition between different regimes in response to a graded perturbation. Together with the analysis of the yeast deletion collection, this highlights how the proposed framework serves as a quantitative tool for efficiently scanning different experimental conditions to detect specific defects.

Furthermore, the analysis of mammalian replication dynamics and the microscopic model we introduced both show that how, from DNA content measurements alone, the method can infer DNA replication dynamics with comparable precision to existing alternatives requiring more, costly data. Taken together, our work provides a robust and scalable quantitative approach to infer cell cycle dynamics from flow cytometry data.

Currently RepliFlow relies on some assumptions that may limit its application in specific contexts. First, we have exclusively focused on populations undergoing exponential growth, where the age-distribution across the population is given by Equation 5. The analysis of populations in different regimes, would require quantifying the mode of growth alongside the cell cycle dynamics and deriving the corresponding age-distribution. Second, our framework does not currently account for cell-to-cell variability, so all inferred quantities represent population averages. While our model could incorporate variability (see Materials and Methods), doing so would require additional parameters, increasing complexity and computational cost. Ad-hoc future extensions of RepliFlow can therefore be developed to accommodate specific usage in conditions where our general assumptions are not applicable.

Importantly, the modular nature of our framework can be exploited to develop custom-made models of DNA replication, opening the way to further quantitative investigation of the dynamics of the eukaryotic cell cycle in perturbed or pathological conditions. A promising direction for future work is the development microscopic models of DNA replication for specific perturbations that reproduce the observed DNA profiles where the underlying molecular details are not yet well understood.

Finally, the scalability of the proposed framework allows to quickly scan a large number of experimental conditions. Inspired by research on the constrains underlying proteome allocation [43–45], an important question for future work is whether there are any fundamental constraints that regulate the time allocated to the different cell cycle phases and how these depend on environmental variables.

## MATERIALS AND METHODS

## Appendix A: Derivation of the likelihood function of measuring a distribution of DNA content in an asynchronous population

In this section we introduce a general likelihood-based approach to infer cell cycle dynamics from high-throughput flow cytometry data. This approach not only takes into account single-cell and population level contributions to the DNA content distribution, but also the impact of technical noise. First, we begin by writing the likelihood of measuring an amount of DNA content *y* in a given cell as

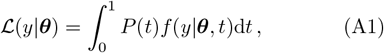

where *t* is the fraction of time elapsed since the start of the cell cycle, *P* (*t*) is the age distribution across the population measured in terms of cell cycle progression, ***θ*** = (***θ***_dyn_, ***θ***_noise_) is a parameter vector controlling the cell cycle dynamics and the technical noise and *f* (*y*| ***θ***, *t*) is the probability of measuring DNA content *y* given the age of the cell *t* and the parameters ***θ***.

As the DNA measurements are independent events, the likelihood of observing a certain DNA distribution ***y*** = {*y*_1_, …, *y*_*N*_} across the population is given by the product of the individual likelihoods

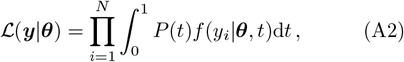

where *N* is the number of cells in the population.

In our model, cell cycle dynamics are fully determined by specifying ***θ***_dyn_, corresponding to the relative duration of the different cell cycle phases and the dynamics of DNA content during DNA replication. Therefore, we represent the dynamics by the parameters ***θ***^*\**^ that maximise the likelihood function

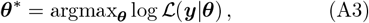

where for convenience we maximise the logarithm of the likelihood function. The maximum likelihood estimator (MLE) is the parameter combination that maximises the probability of observing the empirical data given the model.

To obtain a closed expression for the likelihood function (Equation A2) we have to specify *P* (*t*), the age distribution across the population, and *f* (*y*| ***θ***, *t*), the dynamics of DNA content along the cell cycle. First, we focus on the dynamics of DNA content along the cell cycle. We assumme that every cell follows an identical trajectory since the time of birth (*t* = 0) until the time of division (*t* = 1). Along this trajectory the DNA content of single cells increases monotonically. In particular, DNA content is constant and equal to one copy during G_1_ phase, increases deterministically following some monotonically increasing function *f*_det_(*t*) during S phase, and then remains constant at two copies until the time of division. Additionally, the measured amount of DNA is affected by noise introduced during the measurement process.

We model the measurement noise as a t-Student distribution with location parameter *µ* = 0, scale parameter proportional to the intensity of the signal *τ* = *σf*_det_ ([10, 46, 47]) and degrees of freedom parametrized by *?*. The robustness of the t-Student distribution against the presence of outliers in the data ([48]) makes it a convenient choice to minimise the impact of outliers that are not removed by the preprocessing steps. Taken together, the probability of measuring DNA content *y* given parameters ***θ*** and the cell age *t* reads

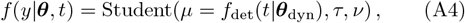

where *f*_det_ represents the deterministic DNA replication dynamics.

At the population level, we consider populations in exponential growth. In that case, the distribution of cell cycle age across the population also follows an exponential form ([13])

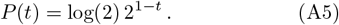

Taken together, the model is defined by Equation A2 together with Equation A4 and Equation A5. The maximisation of the log-likelihood function log with respect to the parameters *θ* is implemented using CmdStanR 2.32.2 in R 4.4.1.

To quantify goodness of fit, we define a fitting score as *s* = 1 − ⟨ *D*⟩, where *D* = sup*x* |*F*_data_(*x*) − *F*_fit_(*x*) | is the two-sample Kolmogorov-Smirnov statistic quantifying the distance between the empirical and inferred CDFs and the average is taken over 100 samples of size *N*. Unless specified otherwise, we choose *s*^*\**^ = 0.95 as a lower threshold for goodness of fit.

Throughout this manuscript we consider cell populations to be homogeneous and described by a parameter vector ***θ***_dyn_. Any differences between cells are attributed to technical noise or their specific positions along the cell cycle. For the sake of completeness, here we briefly describe how to analyse populations where the dynamical parameters ***θ***_dyn_.

Assumming the parameters to be distributed across the population according to a probability distribution *P* (***θ***_dyn_) the likelihood function reads

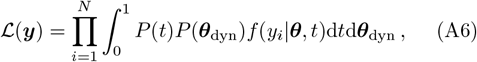

where we have assummed that the heterogeneity is only in the dynamical parameters and not in the parameters quantifying the technical noise.

### Appendix B: Data processing

#### 1. Gating of raw .fcs files

Raw .fcs files were gated using a custom-made Python script. Gating was automated based on the density of events in the forward (FSC-A) and side (SSC-A) scatter channels by binning and thresholding the data in these channels. In addition, events were also gated and reduced on the DNA dye channel in order to eliminate background noise. Finally, events with intensity lower than 10^4^ or larger than 2 · 10^5^ were excluded from further analysis.

The raw .fcs files for the yeast deletion collection were obtained from the files deposited in FlowRepository by [15].

#### 2. Normalisation of the deletion collection results

Our analysis of the deletion collection shows that mutants cluster in two groups in the space of relative phase durations (Figure 2(a). In addition to the deletion mutants, the deletion collection dataset contains in-plate controls corresponding to WT strains. The existence of control profiles in both clusters shows that they are technical artifacts resulting from batch effects between the different plates.

Therefore, to compare the inference results between different plates, we normalise the fraction of time spent in each cell cycle phase against the in-plate controls. Specifically, we define z-scores for each cell *i* in phase *j* as

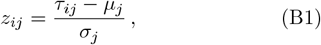

where *τ*_*ij*_ = *t*_*ij*_*/t*_*ij*,C_ is the time spent by cell *i* in phase *j* relative to the in-plate control, *µ*_*j*_ is the average *τ*_*ij*_ across all cells and *σ*_*j*_ is the standard deviation of *τ*_*ij*_ across cells.

#### 3. Calculation of the doubling time of cells

The method infers the relative fraction of time allocated to each cell cycle phase from DNA profiles. To obtain the amount of time that cells spend in each phase we need to independently measure the doubling time of cells *T* in that particular condition.

To obtain doubling times we measured the growth curves of cells in the required conditions. Specifically, we measured the increase in optical density (OD) every 10 minutes from a dilute regime to saturation in media supplemented with 2% glucose. Maximum growth rates *λ* were calculated as the largest slope averaged over a sliding window of 9 time points. Doubling times were calculated from the maximum growth rates as *T* = log(2) *λ*^*−*1^.

### Appendix C: Calculation of the fraction of cells in each cell cycle phase

In this section we describe how we calculate the fraction of cells *ϕ*_*i*_ in phase *i* of the cell cycle starting from the DNA profiles. First, we obtain the fraction of time *t*_*i*_ spent in phase *i* of the cell cycle by fitting the model to the corresponding DNA profile. Then, the fraction of cells in phase *i* of the cell cycle is given by the integral of the age distribution with integration boundaries depending on the fraction of time spent in the respective phases

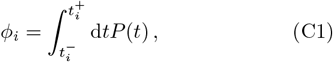

where 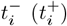 is the fraction of the cell cycle at which phase *i* begins (ends).

Specifically, the fraction of cells in G_1_ reads

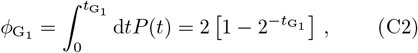

where *P* (*t*) is given by Equation A5.

In the main text, we chose to compare the estimated fraction of cells in G_1_ using our method and the gating approach because its associated region always remains well-defined and identifiable when gating, whereas the definition of the other two regions becomes ambiguous for certain experimental conditions (Supplementary Figure).

### Appendix D: Detailed derivation of the microscopic model

Here we provide a detailed derivation of the microscopic model presented in the main text. The model describes the coupled dynamics of the fraction of replicated DNA, the number of active forks and the number of licensed origins over time. Although we propose a mean field model where we neglect the spatial structure of the DNA, we account for its impact on the dynamics by introducing effective rates of fork annihilation and passive replication, two processes highly dependent on the spatial configuration of the system.

We begin by deriving an equation for the fraction of replicated DNA. DNA is replicated by replication forks travelling along the genome at a rate that is proportional to the number of active forks at time *t*

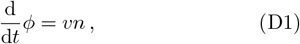

where *v* is an effective replication velocity. Initially the DNA is fully unreplicated *ϕ*(*t* = 0) = 0.

Next, we focus on the dynamics of the number of active forks. Origins fire with rate *γ* producing a couple of replication forks as a result. To take into consideration the completion of replicating regions, we assume that forks can collide with a certain rate. As only forks travelling in opposite directions can meet, the rate of collisions is proportional to the relative velocity between the forks times the probability of randomly choosing two forks travelling in opposite directions times two permutations. Together, the collision rate is 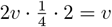. Moreover, as forks can only travel on unreplicated DNA, we take into account the shortening of the unreplicated region as the dynamics progress and modify the collision rate by introducing a factor proportional to the amount of unreplicated DNA 1 − *ϕ*. Taken together, the evolution of the number of active forks reads

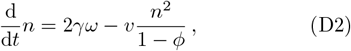

where *ω*(*t*) is the number of licensed origins at time *t*. Additionally, at the start of S phase *n*(*t* = 0) = 0.

For the number of licensed origins, we account for passive replication of origins by active forks. In this case, the rate is proportional to the relative velocity between the fork and the origin, i.e. the replication fork velocity υ. Analogously to the dynamics of the number of active forks, this rate also needs to be modified by a factor proportional to the amount of unreplicated DNA. Therefore, the dynamics of the number of licensed origins read

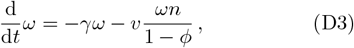

with the initial condition *ω*(*t* = 0) = Ω_0_, where Ω_0_ is the total number of identified budding yeast origins.

The above equations together with the initial conditions fully determine the microscopic model.

To generate the synthetic sequencing time course of Figure 5(e) we run simulations of DNA replication using the inferred microscopic parameters (*v, γ*) together with the confirmed locations of origins in chromosome V of budding yeast (obtained from [49]). The only source of stochasticity in our simulations are the origin firing times, distributed according to an exponential distribution of parameter *γ*. To simulate a population of cells, we average the profiles resulting from 100 realisations of the stochastic dynamics.

### Appendix E: Obtaining microscopic observables from the DNA sequencing data

Synchronised cell populations are a valuable tool to study cell cycle dynamics. In synchronous experiments cells are synchronised at a particular stage of the cell cycle prior to being released and allowed to cycle freely. By sequencing synchronous populations at different times into S phase, we can get insight into DNA replication dynamics at a fine level of detail.

DNA sequencing experiments of synchronous populations return a time series of average copy number *c*_*n*_(*x, t*) for each genomic location *x* at specific time points. Here, we explain how we extract from this data the dynamical observables required to fit the microscopic DNA replication model: the fraction of replicated DNA, the number of active forks and the number of licensed origins.

First, the fraction of replicated DNA is calculated from the average copy number across the genome 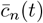 as

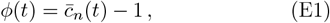

where 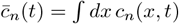 and the integral is performed over the whole genome.

Next, to calculate the number of active forks, we note that replication origins correspond to peaks in the copy number profiles (Figure 5(b)). Therefore, we calculate the number of active forks as twice the number of peaks in the smoothed copy number profiles. To reduce the impact of technical noise, we smooth copy number profiles with a Savietsky-Golay filter prior to peak identification and only identify peaks with a prominence of at least five times the noise level.

The number of licensed origins is calculated as the total number of identified budding yeast origins minus the number of passively replicated and fired origins *ω*(*t*) = Ω_0_ − *ω*_p_(*t*) − *ω*_f_(*t*), where *ω*_p_(*t*) is the cumulative number of origins that have been passively replicated up to time *t, ω*_f_ is the cumulative number of origins that have fired up to time *t* and Ω_0_=829 is the total number of potential origins in *S. cerevisiae* [49]. *ω*_*f*_ is obtained from the copy number profiles as the cumulative number of distinct peaks. Origins located at position *x*_0_ were considered passively replicated at time *t* if *c*_*n*_(*x*_0_, *t*) *>* 1.5 and *x*_0_ did not correspond to a peak of the copy number profile. The locations of potential origins were obtained from OriDB [49].

Finally, each observable was normalised by its maximum value before fitting to the microscopic model.

